# Quinoline synergy and reduced use: a study of pharmacodynamic interactions

**DOI:** 10.1101/2024.09.13.612836

**Authors:** Zahra Sadouki, Emmanuel Q. Wey, Timothy D. McHugh, Frank Kloprogge

## Abstract

**Background:** Meropenem, gentamicin and ciprofloxacin have been used as empiric broad-spectrum combination therapy in different combinations. Recent restrictions on the use of quinolones jeopardises the rational of administering this combination to increase the spectrum of coverage for this particular case. A mechanistic understanding of pharmacodynamic interaction for these combinations is lacking but can provide insight in the necessity of using the different moieties.

**Objectives:** To study pharmacodynamic drug-drug interaction between meropenem, gentamicin and ciprofloxacin against *Escherichia coli*.

**Methods:** Static time kill curve experiments were conducted with *Escherichia coli* (NCTC® 12241) at 0.25 – 16 × MIC for a duration of 24 hours with samples being collected at 0, 2, 4, 6, 8, and 24 hour. Meropenem, gentamicin and ciprofloxacin were tested alone, in two- and three-way combinations. Bacterial load time series data were enumerated on Meuller Hinton plates and Colony Forming Unit data was modelled using nonlinear mixed-effects models in nlmixr.

**Results:** Meropenem, gentamicin and ciprofloxacin two- and three-way combinations prevented regrowth, but did not when these moieties were studied alone. Gentamicin and meropenem were synergistic by decreasing ciprofloxacin IC_50_ and the combination effects of meropenem and gentamicin and the addition of meropenem on top of a gentamicin and ciprofloxacin combination were indifferent.

**Conclusions:** Our findings emphasize the added value of a quinolone in the drug combination. In light of the recent move towards reduced use of quinolones, a quinolone free combination still prevented regrowth, it just did not display further synergy on IC_50_ and was indifferent in initial killing.

## Introduction

It is common practice across healthcare organisations to have local antimicrobial prescribing guidelines and at the Royal Free Hospital London NHS Trust a combination of an aminoglycoside with a beta-lactam or quinolone is recommended for initial empirical treatment of suspected gram negative infections in neutropenic sepsis/sepsis patients *(1)*. In specific clinical syndromes, such as necrotizing fasciitis or fournier’s gangrene all three, i.e. an aminoglycoside, beta-lactam and quinolone, are recommended. Whilst for treatment of other conditions such as bone infections, necrotised devitalised tissue, and biliary sepsis gentamicin may be removed due to inadequate penetration to the site of infection.

The pharmacological rational for administering an antimicrobial drug combination is often based on increasing the spectrum of coverage. In the three-way combination described, meropenem penetrates the bacterial cells by interfering with synthesis of penicillin-binding-protein (PBP). This facilitates increased uptake of ciprofloxacin and gentamicin, and thereby bactericidal activity through prevention of DNA replication and disruption of mRNA translation. However, there is a move towards reducing the use of quinolones, with recent guidance from the MHRA recommending reserving quinolones for cases where other antibiotics are considered inappropriate *(2)*. This reflects concerns over both the side effects of quinolones and the potential for development of resistance.

The aim of this study was to experimentally validate the pharmacodynamic drug-drug interactions between meropenem, gentamicin and ciprofloxacin against *Escherichia coli (E. coli)* through quantitative characterization using a factorial static time-kill approach and nonlinear mixed-effects modelling. The findings enhance our understanding of the mechanism of antimicrobial combination therapy and help inform rationalised use of antimicrobial combination therapy and mitigating the prevalence of Multi Drug Resistant (MDR) gram negative bacterial infections.

## Methods

### Static Time-Kill Experiments

*Escherichia coli* (NCTC® 12241) minimum inhibitory concentrations (MICs) were determined according to CLSI M100-ED32:2022 and were found to be in line with the published CLSI MIC QC ranges for non-fastidious organisms (CLSI Table 5A). 0.015 μg/mL for Ciprofloxacin, 1 μg/mL for Gentamicin and 0.03 μg/mL for Meropenem.

Time-kill assays were conducted following CLSI M26-A guidelines using 96-well plates. Serial dilutions in CAMHB were prepared for meropenem (Sigma, UK), ciprofloxacin TOKU-E, US), and gentamicin (Sigma, UK) monotherapies and combination therapies. Concentrations ranged from ½ to 16 times the MIC, with two-way and three-way combinations adjusted proportionally. Plates were inoculated with 10^5^ CFU/mL *E. coli* and were incubated at 37.5 °C for 24 hours. Bacterial counts (colony forming units, CFU) were performed hourly for the first 8 hours, then at 24 hours on Mueller-Hinton agar, with resistant subpopulations selected on agar supplemented with 2X and 8X MICs of each antibiotic. Data points represent the mean of biological duplicates and technical triplicates.

### Pharmacodynamic Modelling

A nonlinear mixed effects model built in nlmixr 2.0.7 using R 4.1.3 was used to describe the bacterial CFU/mL growth and killing dynamics time series data. Random effects (η) were distinguishing variability between experiments from residual variability (ε) within experimental time series data. Parameters with a biological lower limit of quantification at 0 were log-normally parameterised:

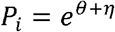

The structural model comprised a logistic growth model to embed biological plausibility.

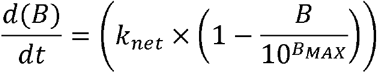

The parameters k_net_, B and B_max_ represented net growth rate, bacterial load and maximum carrying capacity. Drug parameters included maximum antibiotic effect (E_max_), half-maximal antibiotic effect (IC_50_) and a shape factor (γ) to characterise the effects over a concentration range (C_drug_).

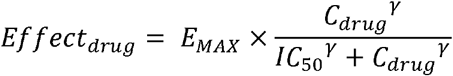

A linear meropenem degradation function was included for all meropenem containing regimens to characterise and embed the chemical degradation occurring at experimental conditions. Drug interation were tested as additivity and Loewe additivity.

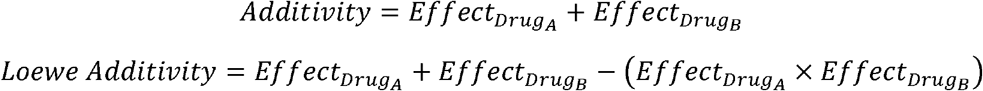

Further synergy was evaluated using a forward stepwise inclusion method using p < 0.05 (ΔOFV = -3.84) for the inclusion of an drug presence, as categorical effect on IC_50_ as additional degree of freedom in a nested model. The ability of models to fit the time series data was further evaluated using goodness of fit plots *(3)*.

## Results

### Static kill curve experiment results

Figure 1 displays the *in-vitro* killing dynamics from time kill experiments for ciprofloxacin, gentamicin and meropenem monotherapy, two-way and three-way combinations against *E. coli* (NCTC® 12241). Data was stratified by MIC to dissect killing dynamics between the different regimens.

**Figure 1:**
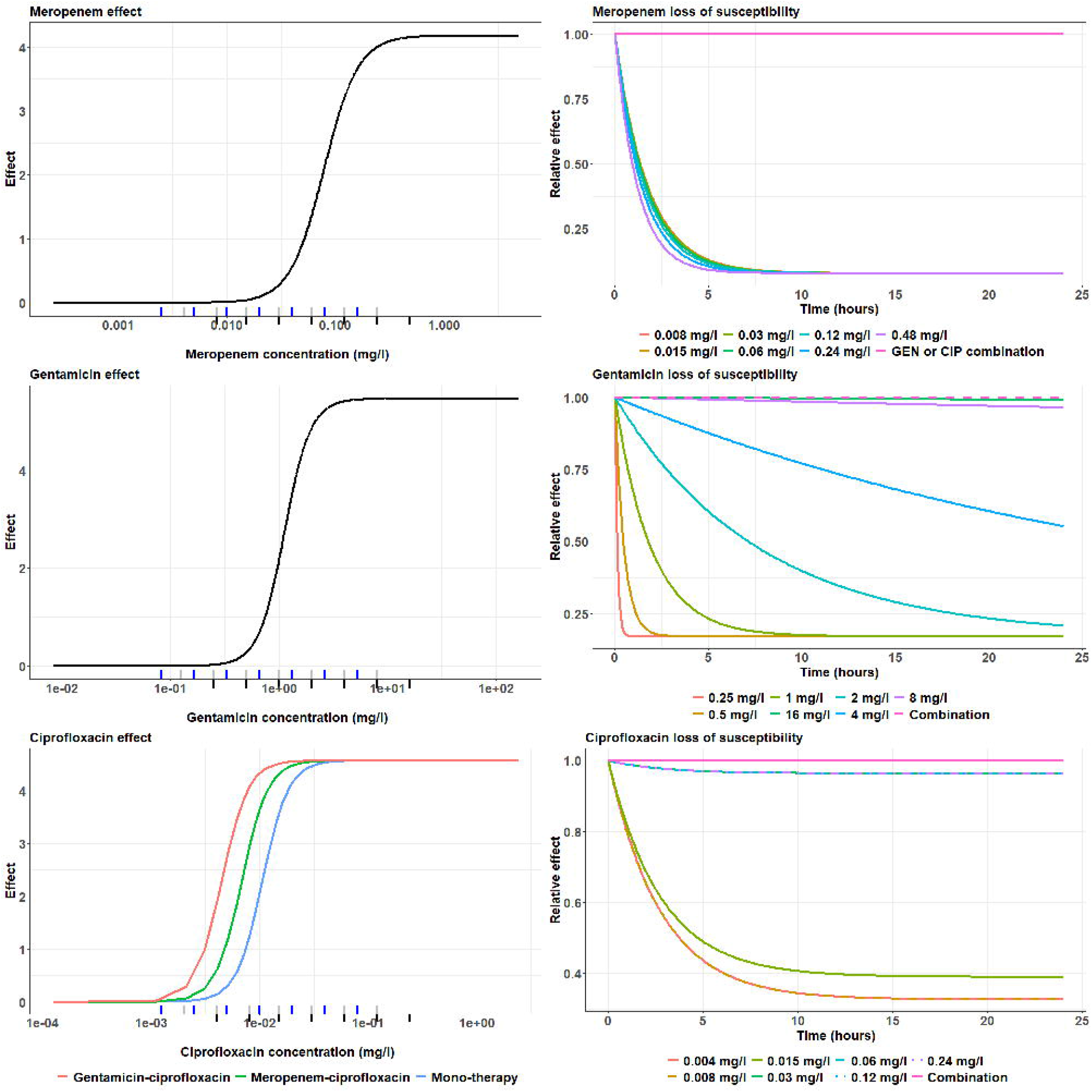
Static time kill curves of meropenem (mer), ciprofloxacin (cip) and gentamicin (gen) antibiotic combinations. CFU/mL over 24 hours of NCTC® 12241 *E. coli* against concentration ranges from 0.25 times MIC to 16 times MIC. Data are stratified by multiplicities of MIC tested (panel) and by combination tested (colour). Biological repeats are plotted independently, and each data point represents the geometric mean of three technical repeats. Antibiotic free growth control represented as 0 X MIC panel (top left).

Different inoculum sizes were studied for the drug free experiments to provide a heterogenous range of growth data (Figure 1). Maximum carrying capacity at 10^11^ CFU/mL was somewhat lower for lower inoculum size experiments, i.e. low and high of 10^3^ and 10^5^ CFU/mL. No difference in killing dynamics was observed at the highest and lowest multiplicity of MIC, i.e. 16 x and 0.25 x MIC. Ciprofloxacin-gentamicin, as two-way combination, displayed the same antimicrobial effect compared to the meropenem-ciprofloxacin-gentamicin, as three-way combination, with all multiplicities, except 0.25 x MIC, producing killing below Limit Of Detection (LOD). The other two-way combinations, i.e. meropenem-gentamicin and ciprofloxacin-meropenem, required ≥ 2 x MIC to achieve comparable killing to ciprofloxacin-gentamicin.

### Quantifying growth and killing

Differences in killing dynamics were investigated in detail by using nonlinear mixed-effects modelling to dissect and quantify pharmacodynamic drug-drug interactions and determine antagonistic and synergistic effects (Suppl. Fig 2 & 3).

Growth control time series data from 24 experiments were fitted using a logistic mixed-effects model with random effects on variability between experiments and residual effect on variability within experiments (Table 1). The effect of inoculum size (i.e. 10^3^ or 10^5^ CFU/ml) on structural model parameter B (inoculum size) and B_MAX_ (maximum carrying capacity) significantly improved the model fit whilst the structural model parameter k_net_ (net bacterial growth) remained unaffected.

**Table 1:**
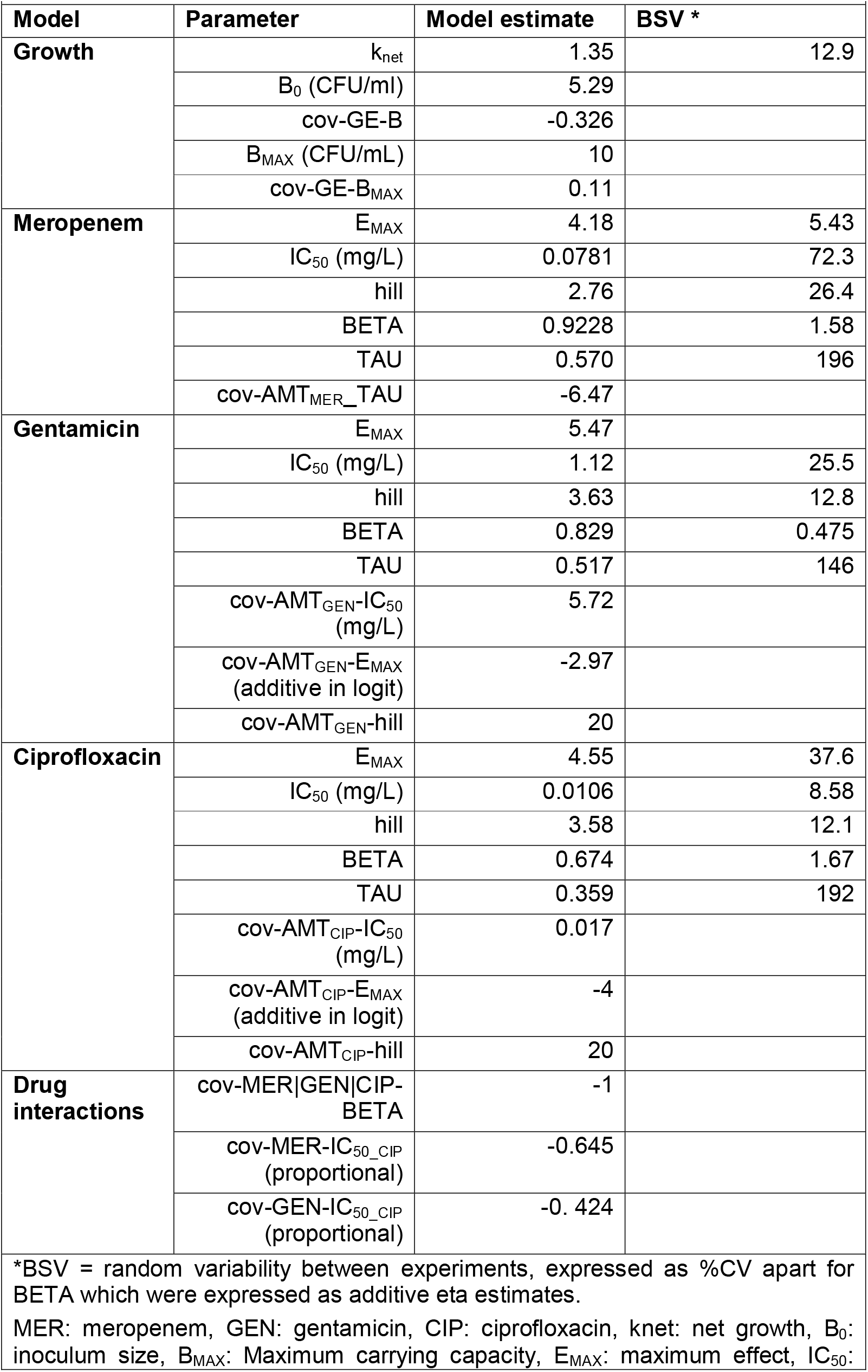
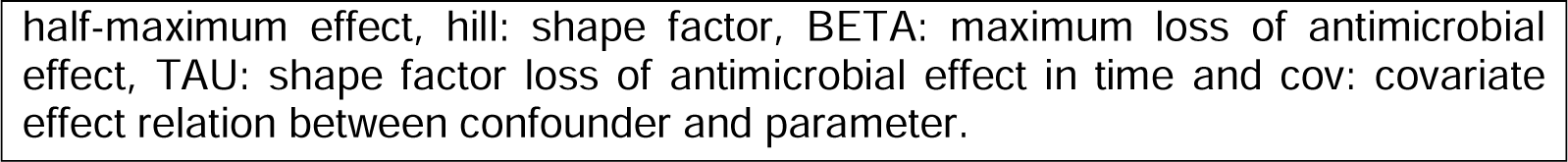
Final parameter estimates of the pharmacodynamic model.

Chemical degradation was accounted for in Meropenem concentration calculations as we found that meropenem concentrations at 37°C dropped by 10% in 8 hours; this degradation was extrapolated for the 24 hour duration of our static kill curve experiment (Suppl. Fig 1). Meropenem E_MAX_ (Maximum effect) was estimated at 4.18 mg/L whilst IC_50_ (concentration causing half-maximum effect) was estimated at 0.0781 mg/L (Table 1) while tested concentrations covered the critical points in the concentration-effect curve (Figure 2). None of the tested meropenem concentrations could reverse inhibition of the killing effect over time, with 16 x MIC (0.48 mg/L) only being able to produce a relative effect of 0.5 at 24 hours compared to at baseline (Figure 1 & 2).

**Figure 2:**
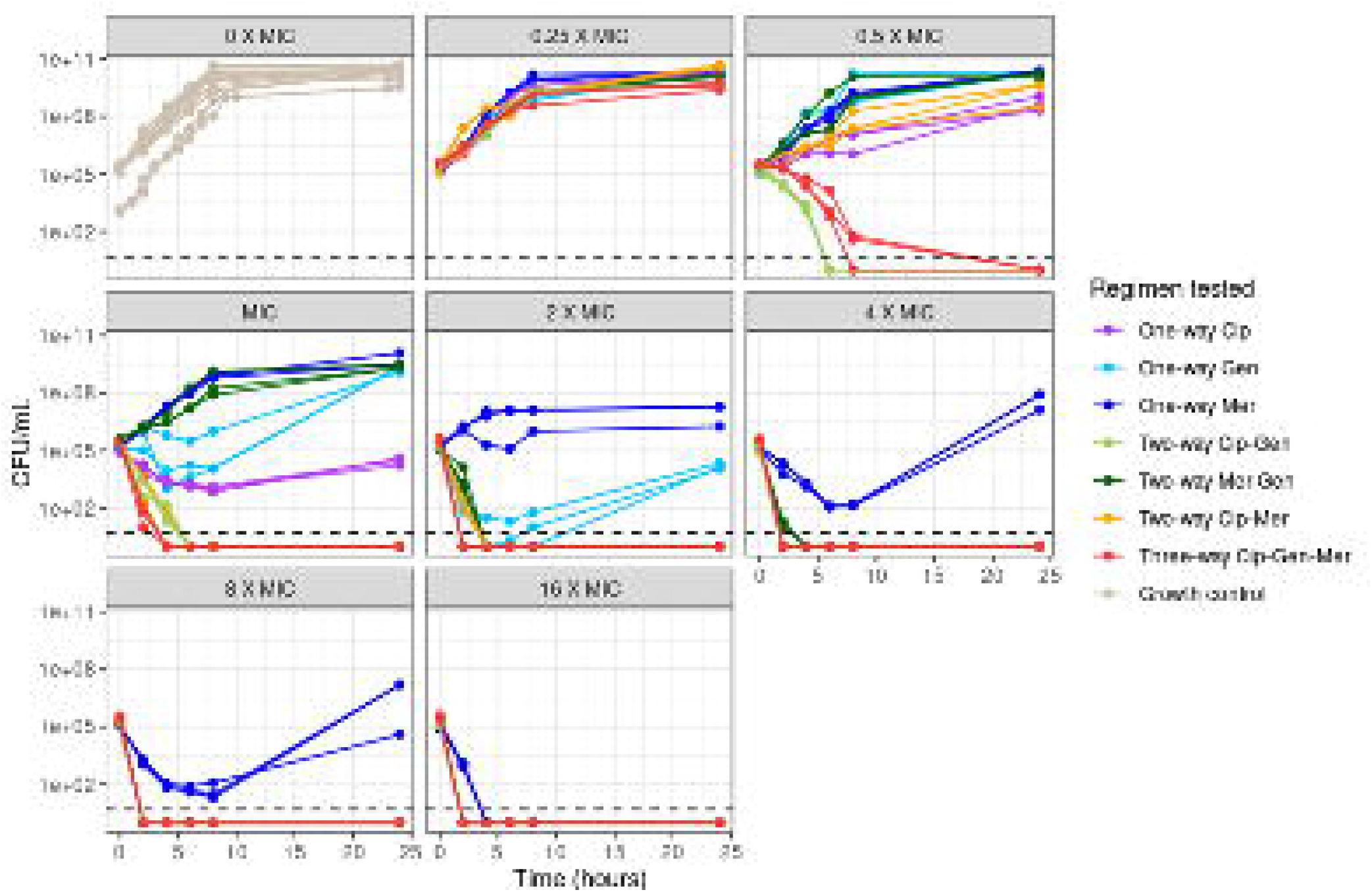
Bacterial killing effect and emergence of bacterial regrowth for meropenem, gentamicin and ciprofloxacin containing regimens. A. Concentration-effect curve to show the killing effect observed (y-axis) against the scaled antibiotic concentration (x-axis). The markings on the x-axis represent the concentrations tested (blue; one-way regimen, grey; two-way regimen and red; three-way regimen) B. Relative effect of the emergence of bacterial regrowth (y-axis) over time (x-axis).

For gentamicin tested concentrations covered the critical points in the concentration-effect curve (Figure 2) and E_MAX_ was estimated at 5.47 mg/L whilst IC_50_ was estimated at 1.12 mg/L (Table 1). Gentamicin killing could be sustained, without presence of waning, for the highest exposures (Figure 1) whilst the equivalent of 8 and 16 x MIC produced a relative effect of 0.5 and near 1 at 24 hours compared to baseline (Figure 2).

Off all three drugs in mono-therapy ciprofloxacin displayed most consistent killing with little presence of a waning effect over time (Figure 1). E_MAX_ (Maximum effect) was estimated at 4.55 mg/L whilst IC_50_ (concentration causing half-maximum effect) was estimated at 0.0106 mg/L (Table 1) and tested concentrations covered the critical points in the concentration-effect curve (Figure 2). Ceasing of ciprofloxacin drug effect did not occur at concentrations > 1 x MIC (Figure 2).

### Quantifying pharmacodynamic drug-drug interactions

All tested drugs in combination, whether in two- or three-way, resulted in sustained killing during the 24-hour experiment without evidence of regrowth, unlike the effect seen with mono-therapy (Table 1 & Figure 1 & 2).

The combined gentamicin-ciprofloxacin effect was additive with a synergistic proportional effect of gentamicin presence on ciprofloxacin IC_50_ -0.424 (Table 1). This resulted in greater activity at 0.5 and 1 x MIC when gentamicin-ciprofloxacin was given in combination compared to alone (Figure 1). Addition of meropenem, i.e. the three-way combination of meropenem-gentamicin-ciprofloxacin, resulted in a similar killing trajectory compared to the two-way gentamicin-ciprofloxacin combination, with the exception of 0.5 and 1 x MIC (Figure 1). Adding meropenem as additive drug effect on top of the gentamicin-ciprofloxacin two-way combination did not yield an improvement of the model fit, indicating the interaction was indifferent.

The combined meropenem-ciprofloxacin effect was additive with a synergistic effect of meropenem presence on ciprofloxacin IC_50_ was also proportional at 0.645 (Table 1). At 1 and 2 x MIC, this yielded antimicrobial effects of meropenem and ciprofloxacin that were greater in combination compared to when they were given alone (Figure 1).

No additional antimicrobial effects were observed when meropenem and gentamicin were given in combination, compared to when gentamicin was given alone. Apart from the 1 x MIC condition, the meropenem-gentamicin combination experiment followed the same killing trajectory up to the point of regrowth for monotherapy.

## Discussion

Here we dissected the role of each antibiotic in the meropenem-gentamicin-ciprofloxacin combination regimen, thus identifying and quantifying the directionality of synergism.

Gentamicin and with cell-wall acting antibiotics have been reported synergistic *(4-7)*, but we only found that the meropenem-gentamicin two-way combination reversed inhibition of the antimicrobial effect over time while no synergy was observed on the cidal effects. Whilst others found up to 5-fold reductions in MIC, with from 64 ug/mL to 2 ug/mL were reported for carbapenem and gentamicin when given in combination *(4)*. Amikacin and meropenem in combination have also be reported synergistic, through a reduced MIC and increased capacity to reduce biofilm formation *(8, 9)*. However, we were conducting time-kill experiments and enabling us to disentangle early cidal effect from regrowth, something that cannot be picked up by MIC experiments only. Given that our results show no regrowth in presence of the meropenem-gentamicin two-way combination our findings in line with previous reports. Moreover, colleagues often reported synergy in resistant bacteria whereas this investigation was conducted using a sensitive laboratory strain *(6, 9-12)*.

Incorporation of Ciprofloxacin in the regimen results in increased pharmacodynamic efficacy as combination with either meropenem or gentamicin reduces the ciprofloxacin’s IC_50_. This corresponds with literature in which meropenem and ciprofloxacin showed synergy against 18/52 Acinetobacter baumannii strains *(12)*. The three-way drug combinations did not display greater antimicrobial effects compared to any of the two-way combinations when challenging the sensitive laboratory stain *E. coli* NCTC 12241.

The inoculum effect we found is in line with other reports. Brook et al. wrote in 1989 that bacterial susceptibility to antibiotics is reduced when the inoculum is increased, thus creating a range of variability of CFU/mL *(13)*. Likewise, meropenem chemical degradation was in line with reports of meropenem being unstable in solution due to degradation of the β-lactam ring. Others reported varying rates of degradation ranging from 90% in 5.7 hours to 20% after 24 hours *(14-16)*.

In conclusion, all two- and three-way combinations of meropenem, gentamicin and ciprofloxacin prevent regrowth as opposed when these moieties were studied on their own. Ciprofloxacin antimicrobial effects improved in combination with either meropenem or gentamicin through reduced ciprofloxacin IC_50_. The combination effects of meropenem and gentamicin were indifferent as well as the addition of meropenem on top of a gentamicin and ciprofloxacin combination.

In light of the recent move towards reduced use of quinolones *(2)* our findings emphasize the added value of a quinolone in the drug combination. On the other hand, a quinolone free combination of meropenem and gentamicin did prevent regrowth, it just did not display further synergy on IC_50_ and was indifferent as opposed to meropenem and gentamicin effects on ciprofloxacin IC_50_ in ciprofloxacin containing combinations.

## Transparency declarations

None to declare

## Author contributions

ZS, EW, TM and FK conceived the project and designed the experiments. ZS conducted the experimental work and FK and ZS analysed the data. ZS and FK wrote the first draft and all others provided feedback.

## Data availability statement

All data is available upon reasonable request.

