## Supplemental data for "Quinoline synergy and reduced use: a study of pharmacodynamic interactions"

Suppl Fig 1: Meropenem degradation over time (hours) for observations (DV) individual predictions (IPRED) and population predictions (PRED)

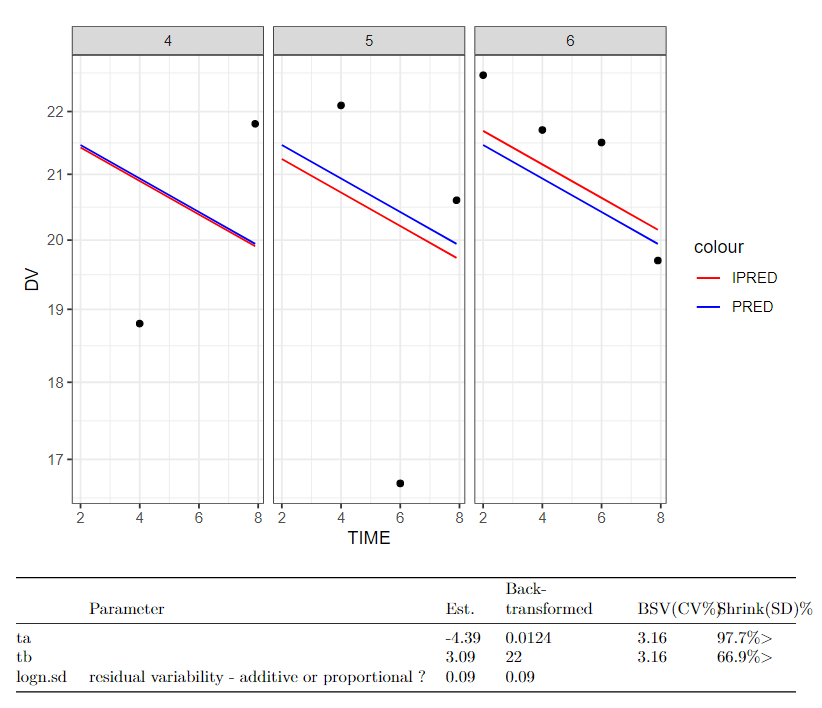

**Suppl Fig 2: Goodness of fit plots displaying observed bacillary load against population predicted (left) and individual predicted (right) bacillary load. The red solid line represent the unity line. Black and red dots represent observed and censored bacillary load samples.**

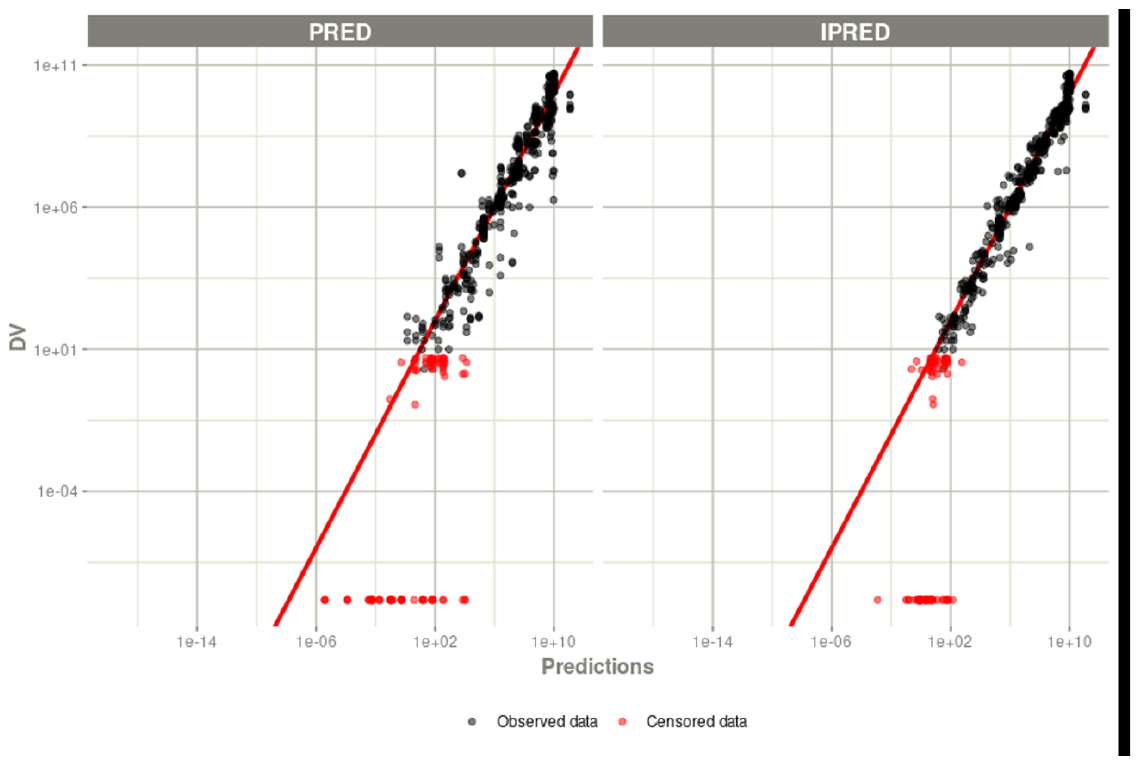

**Suppl Fig 3: Visualisations of mathematical modelling of static time kill data: Model goodness of fit plots. Observed CFU/mL over time (hours) versus the model individual predictions (IPRED) and population predictions (PRED) for each individual experiment.**

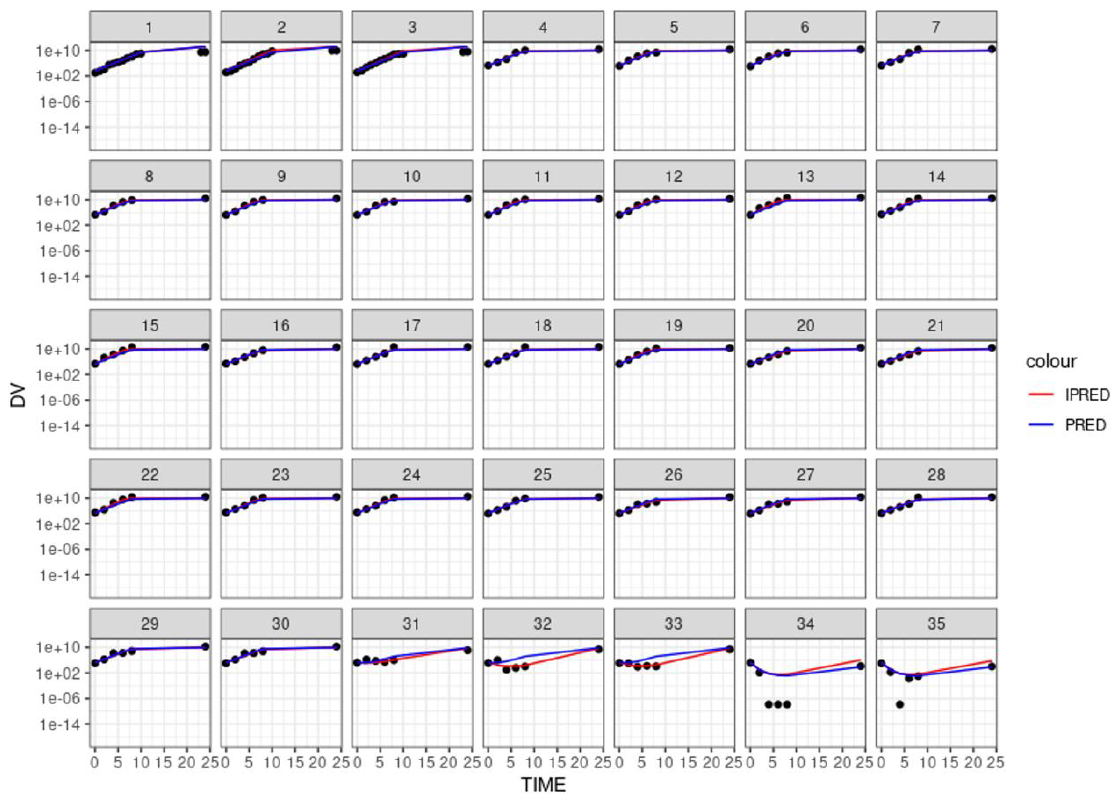

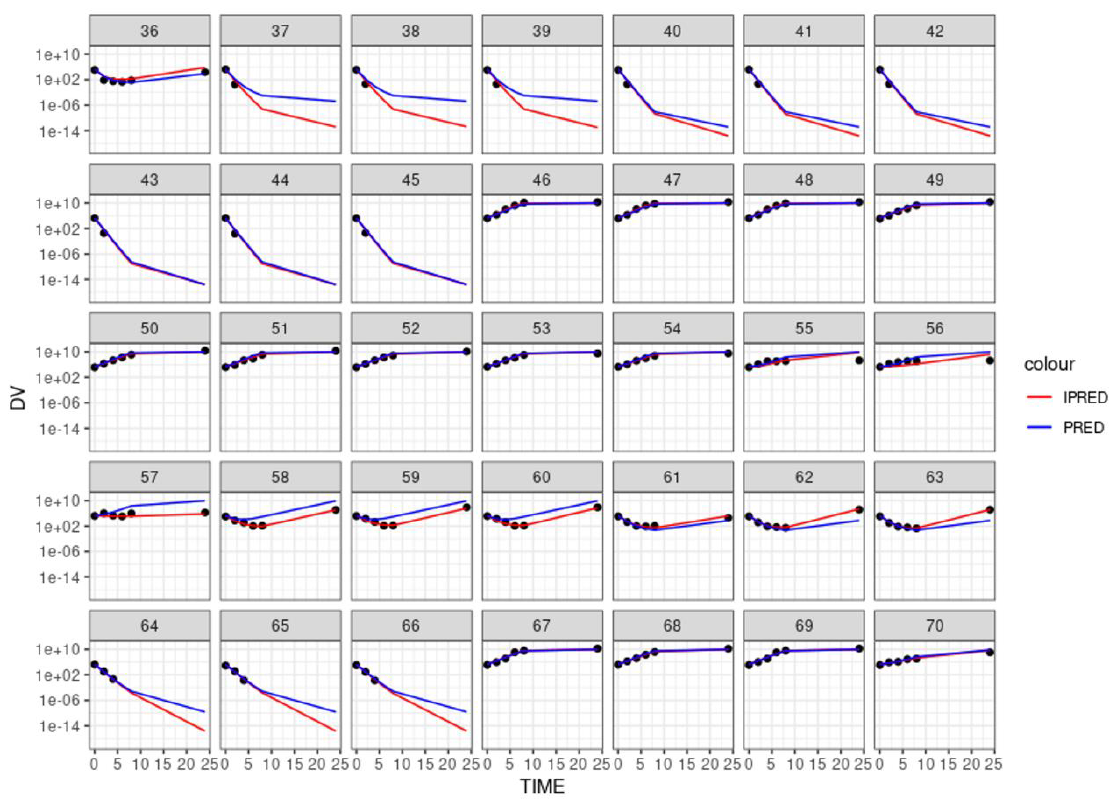

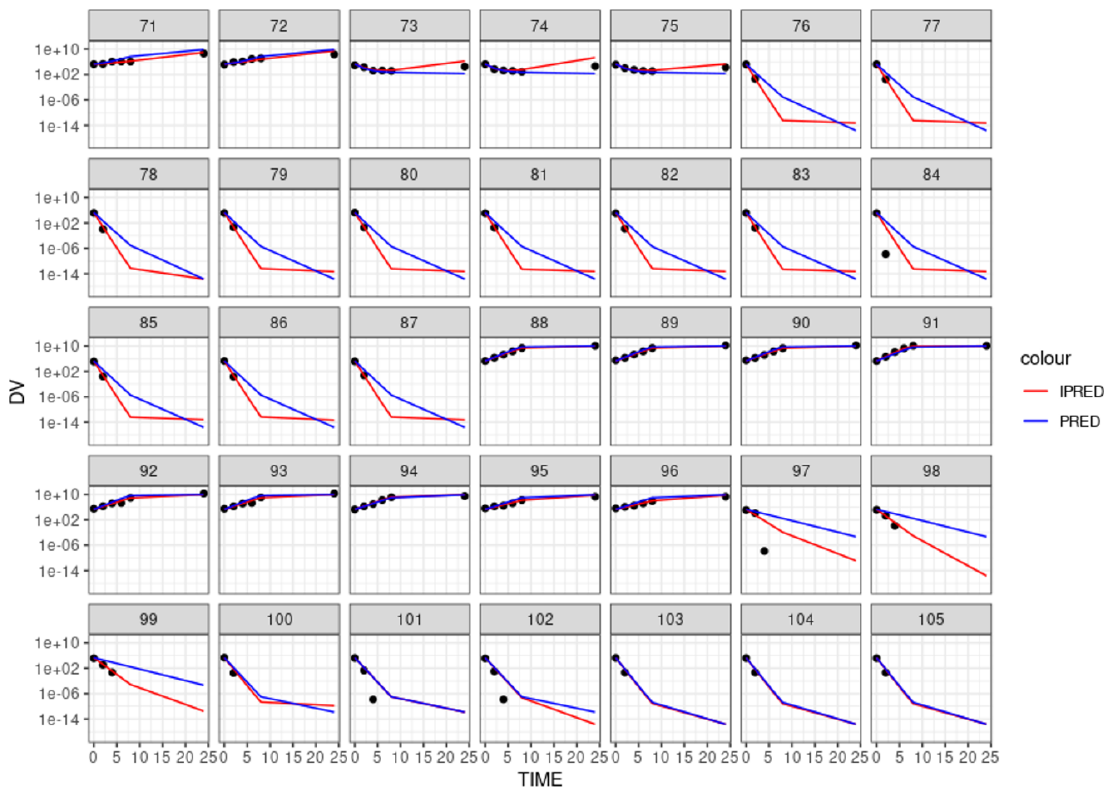

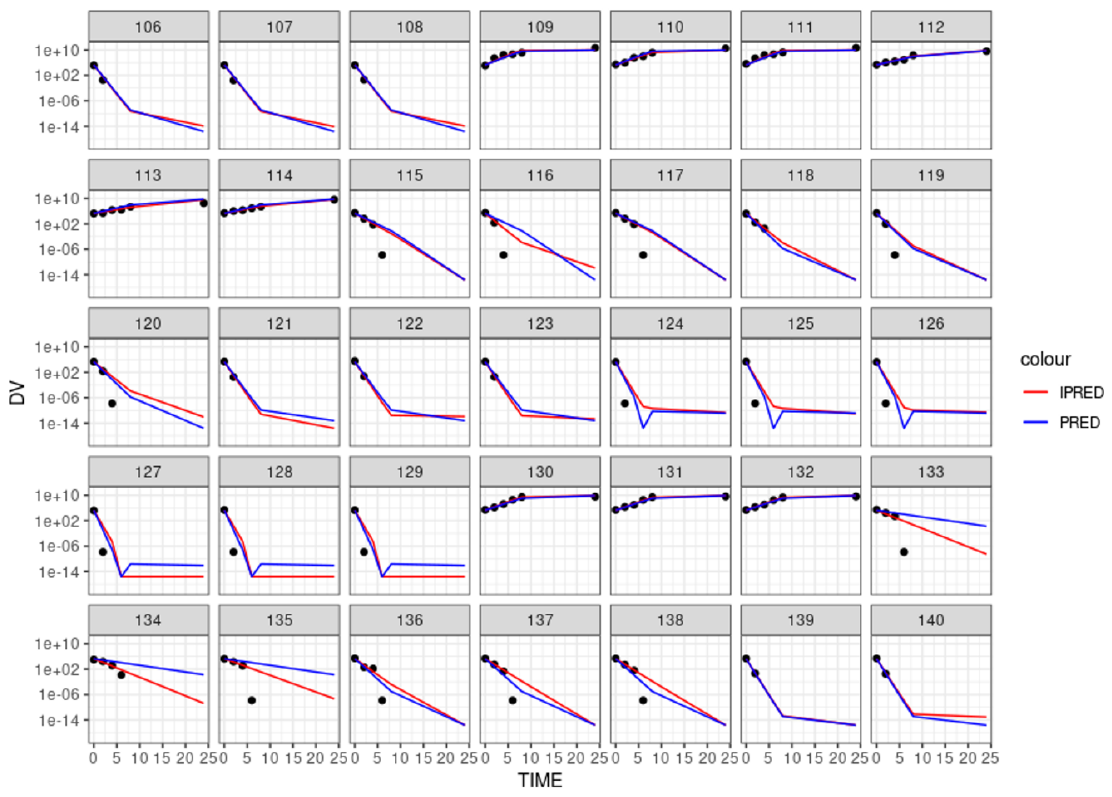

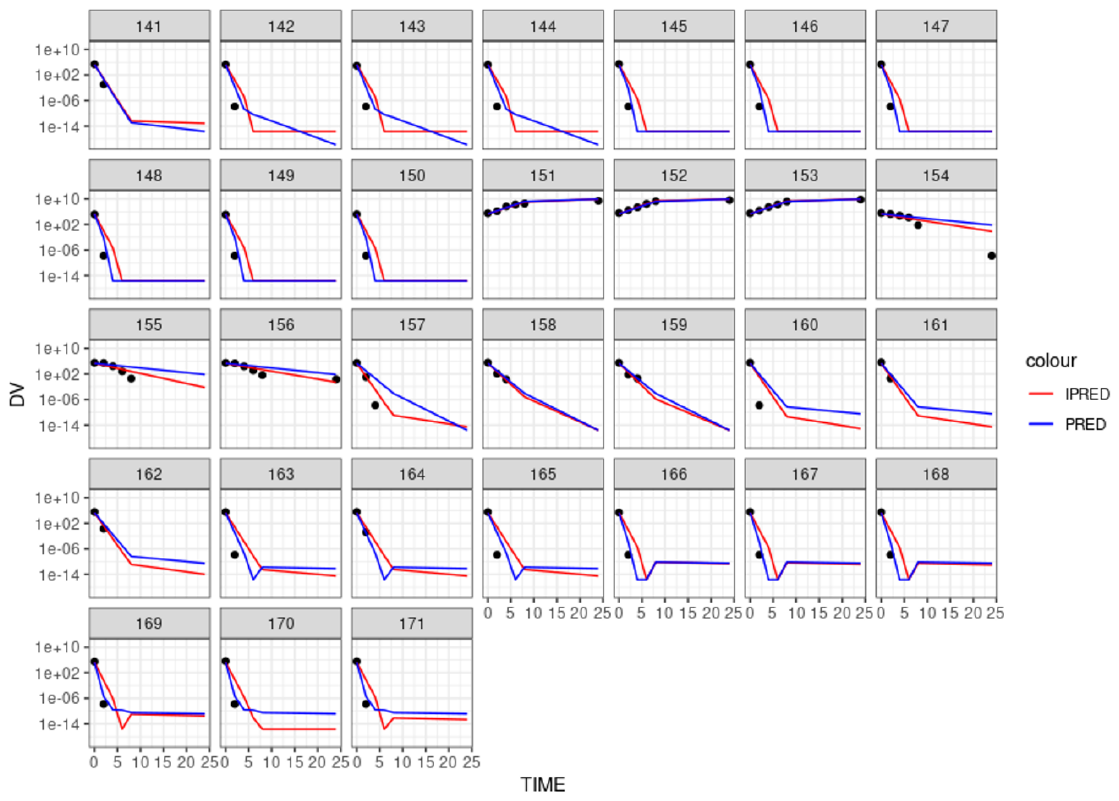
